# Novel personalized cancer vaccine platform based on Bacillus Calmette-Guèrin

**DOI:** 10.1101/2021.03.05.434062

**Authors:** Erkko Ylösmäki, Beatriz Martins, Manlio Fusciello, Sara Feola, Firas Hamdan, Jacopo Chiaro, Leena Ylösmäki, Matthew J. Vaughan, Tapani Viitala, Prasad S. Kulkarni, Vincenzo Cerullo

**Affiliations:** Laboratory of Immunovirotherapy, Drug Research Program, Faculty of Pharmacy, University of Helsinki, Helsinki, Finland; TRIMM, Translational Immunology Research Program, University of Helsinki, Finland; ValoTherapeutics Oy, Helsinki, Finland; Pharmaceutical Biophysics Research Group, Drug Research Program, Faculty of Pharmacy, University of Helsinki, Helsinki, Finland; Serum Institute of India Pvt Ltd, Calcutta, India; iCAN Digital Precision Cancer Medicine Flagship, University of Helsinki; Department of Molecular Medicine and Medical Biotechnology and CEINGE, Naples University 24 Federico II, 80131, Naples, Italy

## Abstract

Intratumoural bacillus Calmette-Guérin (BCG) therapy, one of the earliest immunotherapies, can lead to infiltration of immune cells into a treated tumour. Here, we have developed a novel cancer vaccine platform based on BCG that can direct BCG-induced immune responses against tumour antigens. By physically attaching tumour-specific peptides onto the mycobacterial outer membrane, we were able to induce strong systemic and intratumoural T cell-specific immune responses towards the attached tumour antigens. These therapeutic peptides can be attached to the mycobacterial outer membrane using a cell-penetrating peptide sequence derived from human immunodeficiency virus Tat, N-terminally fused to the tumour-specific peptides. Alternatively, therapeutic peptides can be conjugated with a poly-lysine sequence N-terminally fused to the tumour-specific peptides. Using two mouse models of melanoma and a mouse model of colorectal cancer, we observed that the anti-tumour responses of BCG can be significantly improved by coating the BCG with tumour-specific peptides. In addition, by combining this novel cancer vaccine platform with anti-PD-1 immune checkpoint inhibitor therapy, the number of responders to anti-PD-1 immunotherapy can be significantly increased.

## Introduction

Bacillus Calmette-Guérin (BCG), a live attenuated strain of *Mycobacterium bovis*, is currently the treatment of choice for urothelial carcinoma in situ (CIS) of the bladder ^1, 2^. BCG has also been used previously as an intralesional monotherapy for in-transit melanoma that has resulted, in some studies, in up to 90% regression of BCG-injected lesions and 17% regression of uninjected lesions in immunocompetent patients ^3–5^. In addition, intralesional treatment with BCG has been combined with topical imiquimod (a toll-like receptor 7 agonist) treatment resulting in a complete response rate of 56% ^6, 7^. BCG is an intracellular pathogen that can modulate the tumour microenvironment (TME) by multiple mechanisms including an induction of a massive secretion of chemokines and cytokines that recruit T cells and other immune cells to the TME, as well as by polarization of M2 macrophages towards a more M1-like phenotype ^8,9^. Recently it was shown that BCG treatment led to enhanced activation and reduced exhaustion of tumour-specific T cells, leading to enhanced effector functions and that BCG-induced bladder cancer elimination required tumour-specific CD4^+^ and CD8^+^ T cells, but not T cells specific for BCG antigens ^10^.

Another class of cancer immunotherapy, immune checkpoint inhibitors (ICIs), using antibodies targeting immune checkpoint molecules such as PD-1, PD-L1 and CTLA-4 have demonstrated induction of long-term tumour regression and durable responses in some cancer patients, with response rates of 10-25% in the majority of cancers ^11^. Patients responding to ICI therapy seem to have a pre-existing antitumor immune response with immune cell infiltration into the tumour, which is then enhanced and rendered functional by ICI therapy ^12, 13^. As a consequence, novel combinational therapies that attract tumour-specific CD8^+^ T cells into tumours to increase the number of responders to ICI therapy are much needed.

In order to increase BCG-induced tumour-specific T cell responses and the antitumour effects of BCG therapy, we developed a cancer vaccine platform based on coating BCG bacteria with tumour-specific peptides for redirecting the immune response more towards the tumour instead of the bacteria itself. We then combined this platform with ICI therapy. Intratumoural administration of this cancer vaccine platform, named PeptiBAC (**pepti**de-coated **bac**illus Calmette-Guérin), increased tumour-specific T cell responses. When used in combination with an ICI against programmed death 1 (PD-1), PeptiBAC reduced tumour growth and sensitized tumours to anti-PD-1 ICI therapy by increasing the number of mice responsive to the combination treatment (PeptiBAC in combination with anti-PD-1 ICI). The PeptiBAC platform was also tested in combination with our recently described cancer vaccine platform PeptiCRAd ^14^ (**pepti**de-coated **c**onditionally **r**eplicating **ad**enovirus) using a heterologous prime-boost vaccination strategy ^15^. The heterologous PeptiBAC prime - PeptiCRAd boost vaccination markedly increased tumour-specific T cell immune responses, by directing the immune responses towards the tumour-specific peptides. The elegance of this platform is the introduction of antitumor immunity-inducing peptides non-genetically to the BCG vaccine, which makes this approach highly adaptable and thus suitable for personalized immunotherapeutic approaches that rely on the identification of patient-specific neo-antigens.

## Materials and methods

### Cell lines and reagents

Murine colon carcinoma CT26.wt cell line was purchased from ATCC and was cultured in high glucose RPMI with 10% foetal calf serum (FBS) (Life Technologies), 1% L-glutamine and 1% penicillin/streptomycin. B16F10.9/K1 cell line was kindly provided by Ludovic Martinet (Inserm, France) and was cultured in high glucose DMEM supplemented with 10% FBS, 1% L-glutamine and 1% penicillin/streptomycin. The cell line B16.OVA, a mouse melanoma cell line expressing chicken ovalbumin (OVA), was kindly provided by Prof. Richard Vile (Mayo Clinic, Rochester, MN, USA). B16.OVA cells were cultured in DMEM with 10% FBS (Life Technologies), 1% L-glutamine, 1% penicillin/streptomycin and 5mg/mL of geneticin. Murine dendritic cell line JAWSII was purchased from ATCC and was cultured in alpha minimum essential medium with 20% FBS (Life Technologies), ribonucleosides, deoxyribonucleosides, 4 mM L-glutamine (Life Technologies), 1 mM sodium pyruvate (Life Technologies), and 5 ng/ml murine GM-CSF (PeproTech, USA). Human lung carcinoma A549 cell line was purchased from NIH and was cultured in OptiPRO™ SFM supplemented with 10% FBS (Life Technologies), 1% L-glutamine and 1% penicillin/streptomycin. All cells were cultured at 37 °C/ 5% CO_2_ and were routinely tested for mycoplasma contamination using a commercial detection kit (Lonza).

### Bacteria

Live attenuated bacillus Calmette-Guérin (BCG) vaccines were obtained from various sources. SII BCG (2-8×10^6^ colony forming units [C.F.U]/vial) and SII-ONCO-BCG vaccine (1−19.2×10^8^ C.F.U/vial), were kindly provided by the Serum Institute of India (Pune, India). BCG vaccine (1.5−6.0×10^6^ C.F.U/vial) was purchased from InterVax (Toronto, Canada), while BCG vaccine AJV (2-8×10^6^ C.F.U/vial) from AJ Vaccines (Copenhagen, Denmark) was a kind gift from Professor Helen McShane (University of Oxford).

### Viruses

An adenovirus expressing murine OX40L and CD40L (VALO-mD901) was used in heterologous prime-boost experiments. The development of VALO-mD901 has previously been described (in press). Briefly, a part of the E3B-region of a pAd5/3-D24 backbone plasmid was replaced with human cytomegalovirus (CMV) promoter region, murine OX40L, a 2A self-cleaving peptide sequence, murine CD40L gene and rabbit β-globin polyadenylation signal. The virus was amplified in A549 cells and purified on double caesium chloride gradients and stored below-60°C in A195 adenoviral storage buffer ^16^. The viral particle (VP) concentration was measured at 260/280 nm and infectious units (IU) were determined by immunocytochemistry (ICC) by staining the hexon protein on A549-infected cells.

### Peptides

The following peptides were used in this study: GRKKRRQRRRPQRWEKISIINFEKL, RWEKISIINFEKL, KKKKKK-SIINFEKL and SIINFEKL (containing an MHC class I-restricted epitope from chicken ovalbumin, OVA_257-264_), KKKKKK-SVYDFFVWL and SVYDFFVWL (containing an MHC class I-restricted epitope from tyrosinase-related protein 2, Trp2_180–188_), KKKKKK-SPSYAYHQF and SPSYAYHQF (containing a modified MHC class I-restricted epitope from murine leukaemia virus envelope glycoprotein 70 [gp70_423–431_] where V5A change was made to the original AH1 epitope for enhanced immunogenicity ^17^). All peptides were purchased from Zhejiang Ontores Biotechnologies (Zhejiang, China).

### PeptiBAC complex formation

0.75×10^5^−12×10^7^ C.F.U of BCG resuspended in PBS were complexed with 40-90 nmol of CPP or polyK-extended peptides resuspended in DMSO and incubated for 15 minutes at room temperature (RT). After complexation, PeptiBAC complexes were pelleted by centrifugation at 1000g for 10 min at RT and the buffer was changed to remove unbound peptides.

### PeptiCRAd complex formation

PeptiCRAd complexes were prepared by mixing VALO-mD901 adenovirus (in A195 storage buffer) with polyK-extended Trp2 epitope (in 0.9% saline) at a ratio of 1.8×10^5^ peptides per one virus particle. The mixture was then incubated at room temperature for 15 min. For animal injections, the complexes were diluted further in 0.9% saline to administration volume.

### Surface plasmon resonance

Measurements were performed using a multi-parametric SPR Navi™ 220A instrument (Bionavis Ltd, Tampere, Finland). Phosphate buffered saline (PBS) (pH 7.4) was used as a running buffer. A constant flow rate of 20 μL/min was used throughout the experiments, and temperature was set to +20°C. Laser light with a wavelength of 670 nm was used for surface plasmon excitation. An Au-SiO_2_ sensor slide was activated by 5 min of plasma treatment followed by coating with APTES ((3-aminopropyl) triethoxysilane) by incubating the sensor in 50 mM APTES in isopropanol for 4 h. The sensor was then washed and placed into the SPR device. BCG was immobilized *in situ* on the sensor surface in two of four test channels by injecting approximately 1-4×10^6^ C.F.U of BCG in PBS (pH 7.4) for 12 min, followed by a 3-min wash with PBS. For testing the interaction between various peptides and the mycobacterial outer membrane, 100 μM of the tested peptides extended with CPP or polylysine sequences, or without the attachment moieties (as non-interacting controls) were injected into a BCG coated channel and into an uncoated channel of the flow cell.

### Cross-presentation experiments

JAWSII cells were seeded in 24 well plates (5×10^5^cells/well) and pulsed with PeptiBAC prepared as previously described by complexing 1.5-6×10^6^ C.F.U of BCG with 40nmol of CPP-OVA peptide (GRKKRRQRRRPQRWEKISIINFEKL) or no peptides. After 24h, cells were collected by scraping and stained with APC-conjugated anti-mouse H-2K^b^ bound to SIINFEKL (141606, BioLegend), PerCP-conjugated anti-mouse CD86 (105025, BioLegend) and FITC-conjugated anti-mouse CD40 (124607, BioLegend) antibodies and analysed by flow cytometry.

### Animal experiments

All animal experiments were reviewed and approved by the Experimental Animal Committee of the University of Helsinki and the Provincial Government of Southern Finland (license number ESAVI/11895/2019). Animals were kept in individually ventilated cages under standard conditions (12h light:dark, temperature- and humidity-controlled conditions) and received ad libitum access to water and food. Animals were monitored daily for symptoms related to distress and pain including hunched posture, overall activity/ability to move and roughness of the hair coat. Tumour dimensions were measured by calliper (largest tumour diameter and perpendicular tumour diameter) every second day, starting on the day tumours were first treated. All injections and tumour measurements were performed under isoflurane anaesthesia.

For the B16-OVA melanoma experiment, 8-to 9-week-old immuno-competent female C57BL/6JOlaHsd mice were injected in the right flank with 350,000 B16.OVA melanoma cells, and were treated 12-, 15- and 22-days post tumour implantation with 0.75 − 3×10^5^ C.F.U/dose of BCG alone, 0.75 − 3×10^5^ C.F.U/dose of PeptiBAC-OVA, peptides alone or PBS as a mock-treated group. On day 27 post tumour implantation, 3 mice from each group were sacrificed and spleens and tumours were collected for ELISPOT and flow cytometry analysis. The remaining animals were followed up for survival.

For the B16F10.9/K1 melanoma experiment, 8-to 9-week-old immuno-competent female C57BL/6JOlaHsd mice were injected in the right flank with 300,000 B16F10.9/K1 cells together with a 1:1 ratio of Matrigel Basement Membrane Matrix High Concentration (Corning, USA), and were treated 8-, 10-, and 22-days post tumour implantation with 6.25×10^6^−12×10^7^ C.F.U/dose of BCG, 6.25×10^6^−12×10^7^ C.F.U/dose of PeptiBAC-Trp2 or PBS as a mock-treated group. Groups receiving anti-PD-1 (InVivoMab, USA, clone RMP1-14) were injected intraperitoneally three times per week with 100 μg/dose starting at day 16 post tumour implantation.

For the CT26 colon experiment, 8-to 9-week-old immuno-competent female BALB/c mice were injected in the right flank with 600,000 CT26 cells, and were treated 11-, 13-, and 25-days post tumour implantation with 6.25×10^6^−12×10^7^ C.F.U/dose of BCG, 6.25×10^6^−12×10^7^ C.F.U/dose of PeptiBAC-AH1 or PBS as a mock-treated group. Groups receiving anti-PD-1 (InVivoMab, USA, clone RMP1-14) were injected intraperitoneally three times per week with 100 μg/dose starting at day 17 post tumour implantation.

For the prime-boost vaccination experiments, 8-to 9-week-old immuno-competent naïve female C57BL/6JOlaHsd mice were treated subcutaneously with 1×10^9^ VP/dose of PeptiCRAd VALO-mD901-Trp2, PeptiCRAd VALO-mD901-OVA, 2-8×10^6^ C.F.U/dose of PeptiBAC-Trp2, 2-8×10^6^ C.F.U/dose of PeptiBAC-OVA or saline as a mock-treated group. Vaccinations were performed 14 days apart. 4 days after the last injection, mice were sacrificed and spleens were collected for ELISPOT assay. All mice strains were obtained from Envigo (Venray, the Netherlands).

### Flow Cytometry

The following antibodies were used in the experiments: TruStain FcX™ anti-mouse CD16/32 (101320, BioLegend), FITC anti-mouse CD8 (A502-3B-E, ProImmune), Phycoerythrin (PE) anti-mouse CD3e (550353, BD Pharmingen), Peridinin-Chlorophyll-Protein (PerCP) anti-mouse CD19 (115531, BioLegend) and PE-Cyanine 7 anti-mouse CD4 (25-0041-82 eBioscience). SIINFEKL epitope-specific T cells were studied using APC-labelled H-2Kb/SIINFEKL pentamer (F093-84B-E, ProImmune). SVYDFFVWL (Trp2) epitope-specific T cells were studied using PE-labelled H-2Kb/SVYDFFVWL pentamer (F185-82B-E, Proimmune). SPSYVYHQF (AH1) epitope-specific T cells were studied using PE-labelled H-2Ld/SPSYVYHQF pentamer (F398-82A-E, Proimmune). Flow cytometric analysis were performed using a BD Accuri 6C Plus (BD Biosciences) or a BD LSRFortessa™ (BD Biosciences) flow cytometer and FlowJo software v10 (BD Biosciences) was used for data analysis.

### Enzyme-linked immunospot (ELISPOT) assays

The amount of SIINFEKL (OVA_257-264_), SVYDFFVWL (TRP2_180-188_), BCG and adenovirus - specific activated, interferon-γ secreting T cells were measured by ELISPOT assay (CTL, Ohio USA) according to the manufacturer’s instructions. Briefly, 2 μg of SIINFEKL or SVYDFFVWL peptide was used to stimulate the antigen presenting cells. After 2 or 3 days of stimulation, plates where stained and sent to CTL-Europe GmbH for counting of the spots.

### Statistical analysis

Statistical analysis was performed using GraphPad Prism 8.0 software (GraphPad Software, USA). For data analysis, one-way ANOVA was used. All results are expressed as the mean ± SEM.

## Results

### Bacillus Calmette-Guérin can be coated with therapeutic peptides by using a cell penetrating peptide sequence or a poly-lysine sequence as an anchor

The mycobacterial cell wall is a highly complex structure containing multiple layers of different lipid components and has an extremely negative surface potential ^18–20^. We hypothesized that therapeutic peptide sequences could be attached into the mycobacterial cell wall using a cell penetrating peptide (CPP) sequence or a poly-lysine sequence as attachment moieties (see Figure 1 for schematic presentation of the PeptiBAC platform). Various CPP sequences were tested by surface plasmon resonance (SPR) for their efficacy at anchoring therapeutic peptides into the mycobacterial cell wall (data not shown), and a CPP sequence derived from HIV Tat protein was found to be the most efficient CPP sequence for anchoring the peptides (Figure 2A). In addition to the CPP sequence derived from HIV Tat, a positively charged poly-lysine sequence was found to efficiently anchor the peptides into the cell wall (Figure 2B and C). We estimated the number of peptides bound to BCG bacterium using these two different attachment moieties. For the SIINFEKL antigen containing an N-terminal CPP Tat sequence, the number of peptides bound to BCG was estimated to be 1.8×10^6^ peptide molecules/bacterium. For the Trp2 antigen and for the AH1 antigen containing N-terminal poly-lysine sequences, the number of peptides bound to BCG was estimated to be 4.4×10^6^ peptide molecules/bacterium and 3.2×10^5^ peptide molecules/bacterium, respectively.

**Figure 1.**
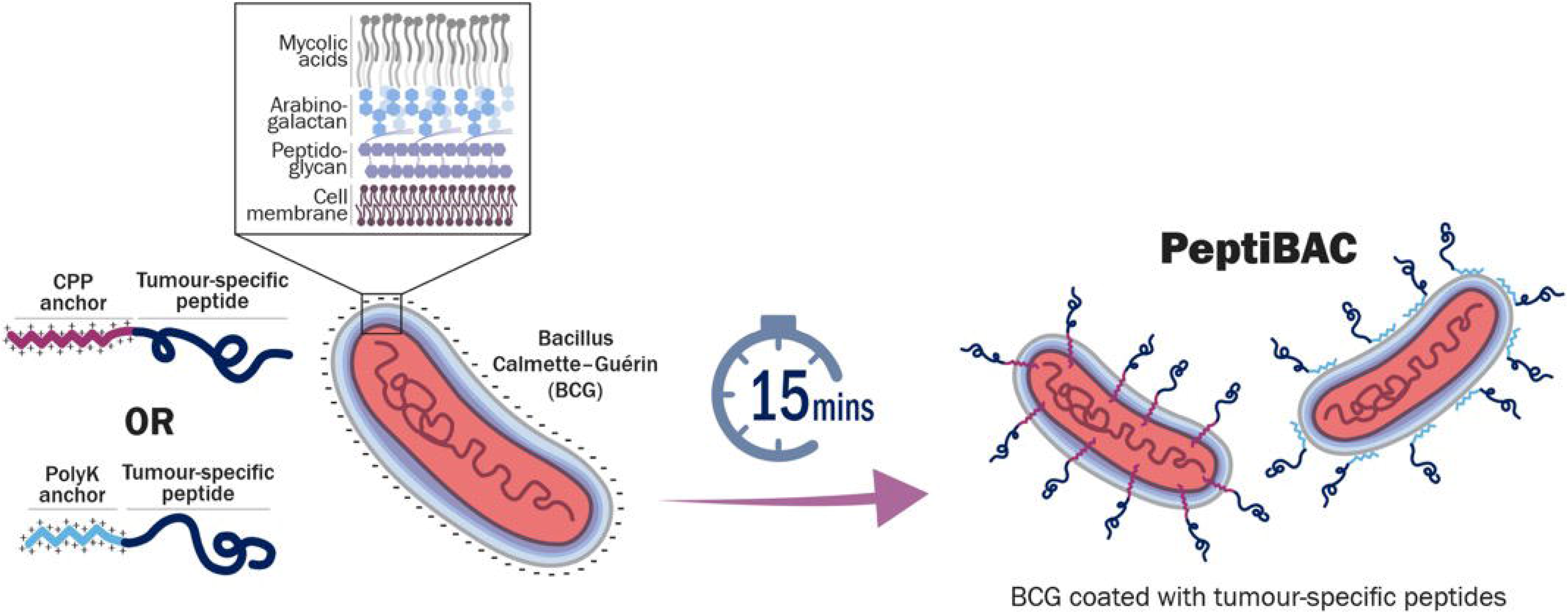
A schematic presentation of a PeptiBAC cancer vaccine platform. Tumour antigens can readily be attached to the mycobacterial outer membrane of Bacillus Calmette-Guèrin (BCG) using a cell penetrating peptide (CPP) sequence or a poly-lysine sequence as an anchoring moiety. Anchor-modified peptides are complexed for 15 min. with BCG for efficient attachment. Unbound peptides are removed by pelleting the bacteria followed by buffer exchange. Various different peptides, including MHC class I and II epitopes, can be delivered by the PeptiBAC-platform for potent activation of antigen-presenting cells and increased antigen-specific immunological responses.

**Figure 2.**
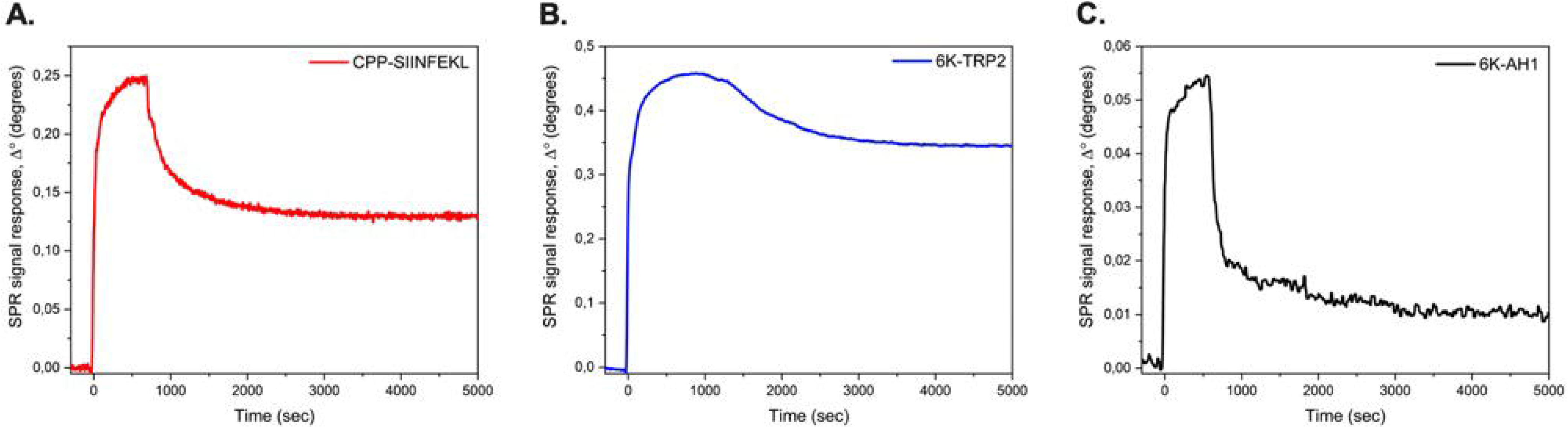
Surface plasmon resonance (SPR) analysis of the peptide/BCG interaction. A) Surface plasmon resonance analysis of the interaction between the CPP-OVA and BCG. B) Surface plasmon resonance analysis of the interaction between the polyK-Trp2 and BCG. C) Surface plasmon resonance analysis of the interaction between the polyK-AH1 and BCG.

### Antigen presenting cells can efficiently present therapeutic peptides delivered by PeptiBAC

Next, we tested whether the PeptiBAC platform can deliver therapeutic peptides to antigen presenting cells (APCs) and if the APCs can cross-present the MHC-I epitope portions from these peptides. PeptiBAC-OVA (BCG coated with CPP-containing immunodominant epitope from chicken ovalbumin; GRKKRRQRRRPQRWEKISIINFEKL) was used to infect JAWSII murine dendritic cells (DCs) for 24h followed by the assessment of the cross-presentation efficacy of the epitope (SIINFEKL) by flow cytometry (Figure 3A). As expected, PeptiBAC delivered SIINFEKL was efficiently cross-presented by the DCs, as almost 40% of JAWSII cells were shown to cross-present the SIINFEKL epitope. In addition, PeptiBAC-OVA was able to induce enhanced DC activation compared to BCG as assessed by the significantly increased expression of cluster of differentiation 86 and 40 (CD86 and CD40) proteins (Figure 3B and C, respectively).

**Figure 3.**
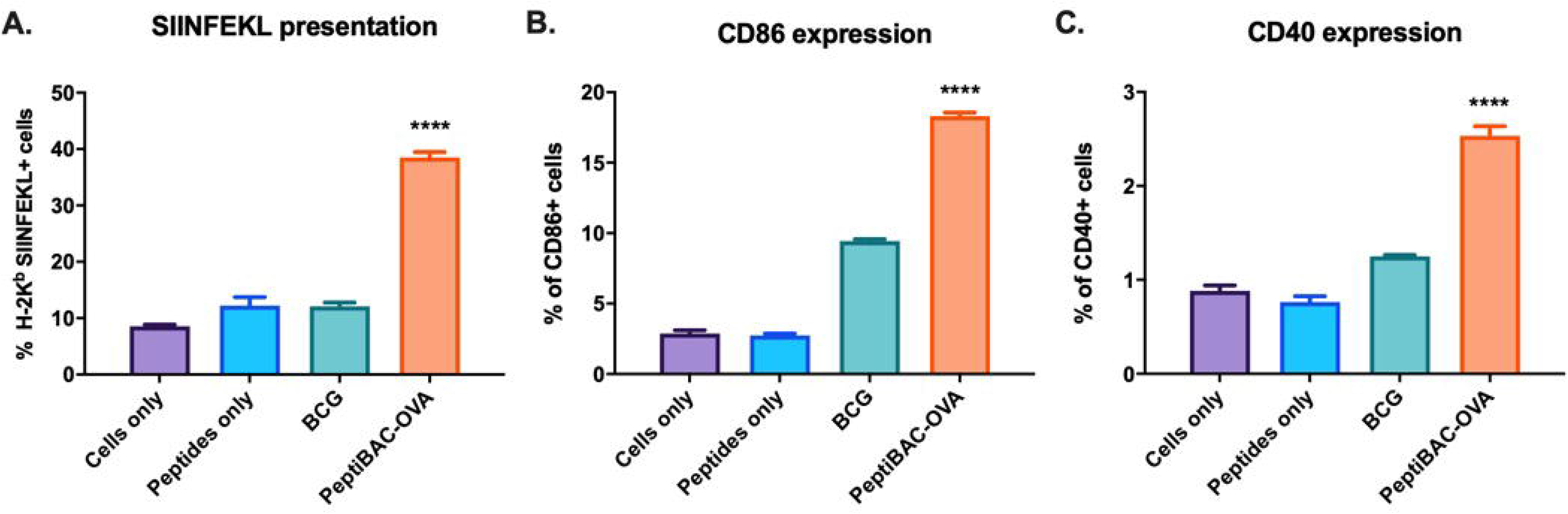
Antigen-presenting cells can readily cross-present antigens delivered by the PeptiBAC platform. Mouse dendritic cell line Jaws II was pulsed with PeptiBAC-OVA, BCG, CPP-containing SIINFEKL peptide alone or left un-pulsed (cells only). Cross-presentation was determined by flow cytometry using APC-conjugated anti-H-2Kb bound to SIINFEKL. CD86 and CD40 expression (as a measure of dendritic cell maturation and activation) was determined by flow cytometry. Each bar is the mean ± SEM of technical triplicates. Statistical analysis was performed with one-way ANOVA. **** p< 0.0001.

### Intratumoural treatment with PeptiBAC with CPP-containing OVA antigen elicits moderate tumour growth control and induces systemic tumour-specific CD8^+^ T cell response in syngeneic mouse model of B16.OVA melanoma

To study the immunostimulatory potential and anti-tumour effects of the PeptiBAC platform, we used a well-established syngeneic mouse melanoma model B16 expressing chicken ovalbumin (OVA) as a model antigen ^21^. When mice bearing B16.OVA tumours were treated intratumourally with OVA-targeting PeptiBAC (PeptiBAC-OVA), BCG, peptides alone or vehicle (mock), we observed a moderate increase in tumour growth control in the PeptiBAC-OVA group as compared to all other treatment groups (Figure 4A). We set a tumour size threshold of 450 mm^3^ for defining the responders in each treatment group. Treating mice with the CPP-containing SIINFEKL peptide alone did not have any significant effect on tumour growth, with one mouse defined as a responder to the therapy in this group. Similarly, in BCG- and mock-treated groups there was one responder in each group; a 13% response rate. In contrast, PeptiBAC-OVA treatment had a moderate effect on tumour growth with three mice defined as responders for the therapy; a 38% response rate for this group of mice. We went on to analyse whether there were any differences in immunological responses against the OVA antigen between the treatment groups, and we first assessed whether there were any differences in the infiltration of immune cells into the tumour microenvironment (TME). We observed that a higher number of cytotoxic CD8^+^ T cells infiltrated into the tumours of PeptiBAC-OVA-treated mice as compared to the tumours of BCG-, peptide alone- or mock-treated mice. However, we did not see any infiltration of tumour-specific CD8^+^ T cell into the tumours in any of the treatment groups (data not shown). In striking contrast to BCG-, peptide alone- and mock-treated mice, a significant induction of a systemic OVA-specific T cell response was seen in PeptiBAC-OVA-treated mice (Figure 4B). The observed moderate increase in tumour growth control in the PeptiBAC-OVA group translated into a clear but non-significant trend towards longer survival, with median survival of 32 days compared to 25, 29 and 27 days with BCG, peptide alone and mock groups, respectively (Figure 4C).

**Figure 4.**
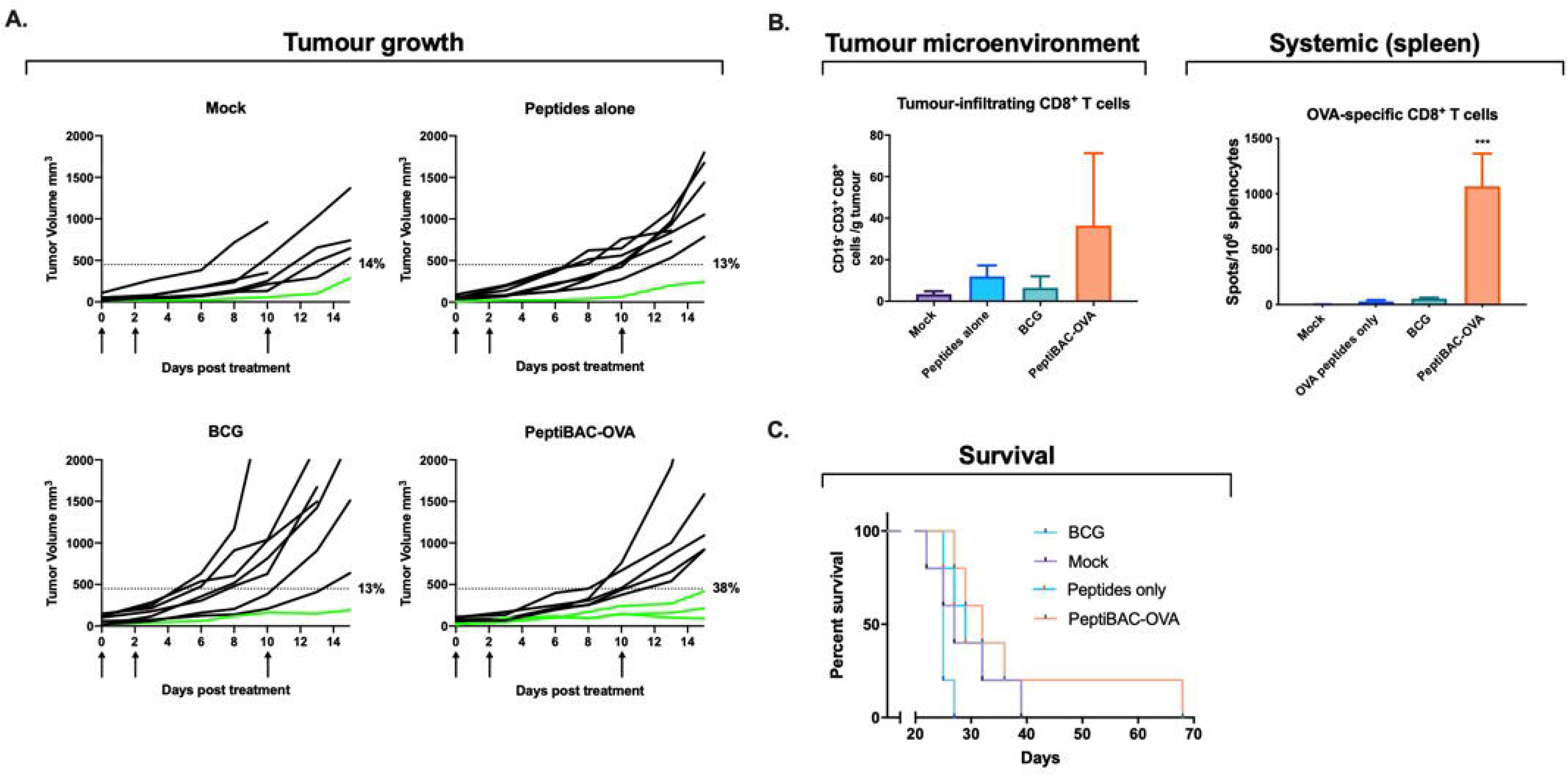
PeptiBAC improves tumour growth control and induces systemic tumour-specific T cell responses in a syngeneic mouse model of B16.OVA melanoma. A) BCG, Peptides alone or PeptiBAC-OVA was given intratumourally 12-, 15- and 22-days post tumour implantation. Individual tumour growth curves for all treatment groups are shown. A threshold of 450 mm^3^ was set to define the percentage of mice responding to the different therapies (dotted line). The percentage of responders in each treatment group is shown on the right side of the dotted line. B) Immunological analysis of tumours and spleens of treated mice. C) Kaplan-Meier survival curve for the treatment groups. The number of mice in each group was 7-8. Statistical analysis was performed with one-way ANOVA. *** p< 0.001.

### Intratumoural treatment with PeptiBAC with poly-lysine-containing Trp2 antigen increases the number of responders to anti-PD-1 therapy, improves tumour control and induces tumour-specific T cell responses in a syngeneic mouse model of B16.F10.9/K1 melanoma

Next, we tested the PeptiBAC platform in a syngeneic mouse model of B16.F10.9/K1 melanoma using a more relevant, tumour-associated antigen from tyrosinase related protein 2 (Trp2_180–188_) in combination with anti-PD-1 immune checkpoint inhibitor therapy (ICI). As the effects of PeptiBAC with CPP-containing OVA antigen on tumour growth control were modest in the previous experiment, we increased the amount of BCG and PeptiBAC given/dose. In addition, we changed the attachment sequence to poly-lysine and tested the effects of combining the PeptiBAC treatment with anti-PD-1 ICI therapy. B16.F10.9/K1 melanoma is a derivative of a highly metastatic B16.F10.9 melanoma with a low cell surface expression of major histocompatibility complex 1 (MHC-I) H-2Kb that was transfected with H-2Kb genes to generate H-2Kb-expressing clone K1 ^22^. The B16.F10.9/K1 clone is more responsive to cancer immunotherapies than the highly immunosuppressive parental strain B16.F10.9. Starting at 8 days post tumour engraftment, mice were treated intratumourally with BCG, anti-PD-1 alone, PeptiBAC-Trp2, BCG in combination with anti-PD-1, PeptiBAC-Trp2 in combination with anti-PD-1 or saline as a mock-treated group. Again, we set the tumour size threshold of 450 mm^3^ for defining the responders in each treatment group. In contrast to mock-treated animals, BCG, anti-PD-1 alone and BCG in combination with anti-PD-1 ICI-treated groups showed modest tumour growth control with response rates of 30%, 27% and 27%, respectively. PeptiBAC-Trp2-treated animals showed robust tumour growth control with a 56% response rate. Remarkably, PeptiBAC-Trp2 in combination with anti-PD-1-treated animals showed efficient tumour growth control, with 70% response rate; increasing the response rate for anti-PD-1 therapy from 27% to 70% (Figure 5A). To further evaluate the mechanism of tumour growth control, we assessed whether there were any differences in the Trp2-specific T cell responses between the treatment groups. We saw increased numbers of tumour-infiltrating CD4^+^ and CD8^+^ T cells in PeptiBAC-Trp2-treated tumours compared to BCG, anti-PD-1 alone and BCG in combination with anti-PD-1 ICI-treated tumours. Also, the number of Trp2-specific CD8^+^ T cells was increased in PeptiBAC-Trp2-treated tumours compared to BCG, anti-PD-1 alone and BCG in combination with anti-PD-1 ICI-treated tumours. In contrast to other treatment groups, PeptiBAC-Trp2 in combination with anti-PD-1-treated tumours had significantly more tumour-infiltrating CD4^+^ and CD8^+^ T cells as well as Trp2-specific CD8^+^ T cells, indicating a synergistic effect on T cell responses by combining the two treatment modalities (Figure 5B, upper panel). We also evaluated systemic tumour-specific T cell responses by analysing the spleens of treated mice. No significant differences in the number of CD4^+^ and CD8^+^ T cells was found between groups. The number of Trp2-specific CD8^+^ T cells was increased in PeptiBAC-Trp2 in combination with anti-PD-1 ICI-treated spleens as compared to other treatment groups, again indicating a synergistic effect on T cell responses by combining the two treatment modalities (Figure 5B, lower panel).

**Figure 5.**
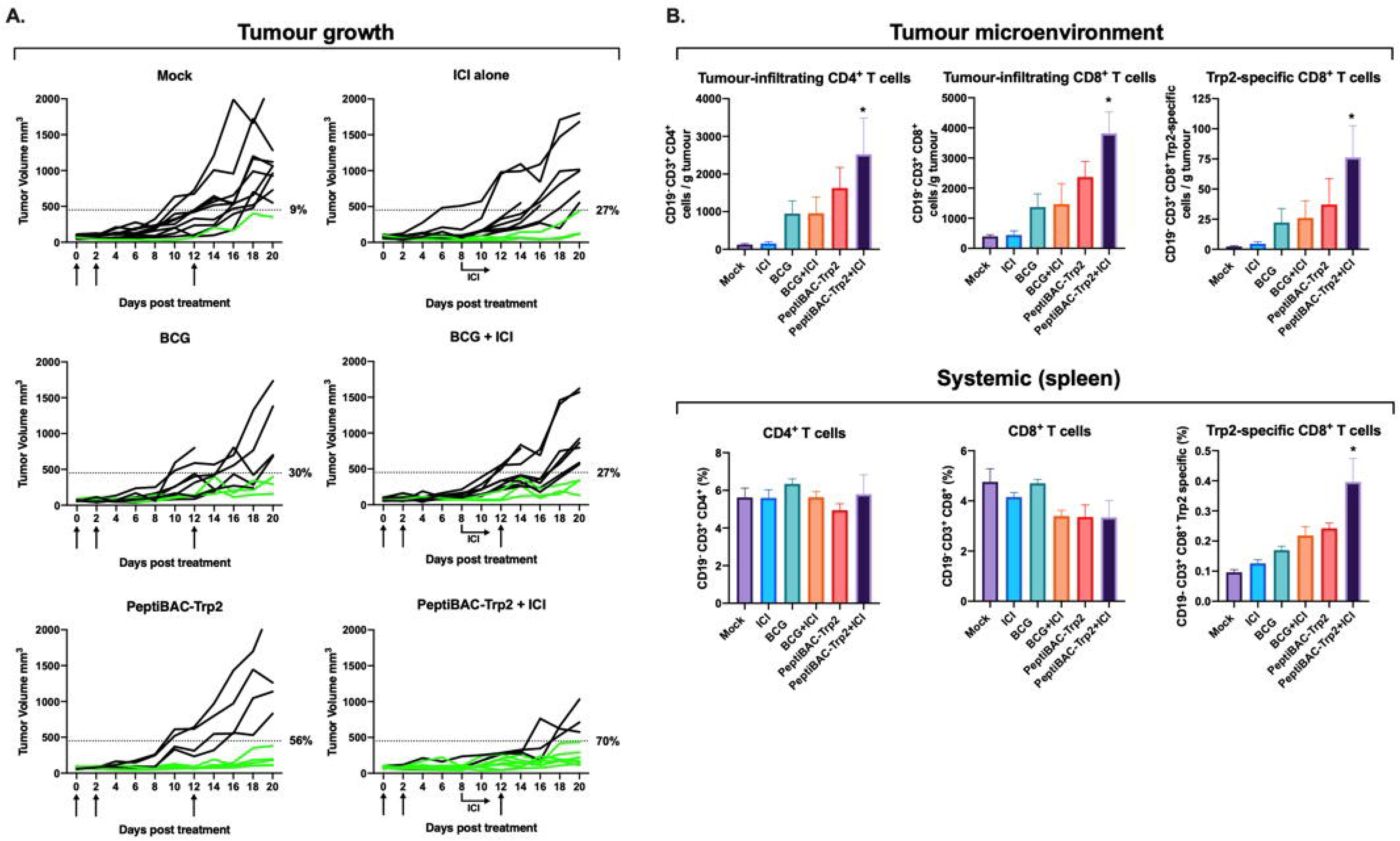
PeptiBAC in combination with anti-PD1 improves tumour growth control compared to either monotherapy and induces robust infiltration of tumour-specific CD8^+^ T cells into tumours in a syngeneic mouse model of B16.F10.9/K1 melanoma. A) Anti-PD-1 immune checkpoint inhibitor alone (100μg/dose given intraperitoneally three times a week, starting at day 8), BCG alone or in combination with anti-PD-1 immune checkpoint inhibitor and PeptiBAC-Trp2 alone or in combination with anti-PD-1 immune checkpoint inhibitor was given intratumourally 8-, 10-, and 22-days post tumour implantation. Individual tumour growth curves for all treatment groups are shown. A threshold of 450 mm^3^ was set to define the percentage of mice responding to the different therapies (dotted line). The percentage of responders in each treatment group is shown on the right side of the dotted line. B) Immunological analysis of tumours and spleens of treated mice. The number of mice in each group was 9-11. Statistical analysis was performed with one-way ANOVA. * p< 0.05.

### Intratumoural treatment with PeptiBAC with poly-lysine-containing modified gp70 antigen increases the number of responders to anti-PD-1 therapy, improves tumour control and induces tumour-specific T cell responses in a syngeneic mouse model of CT26 colorectal cancer

To validate the PeptiBAC platform as a more universal cancer vaccine platform, we tested the platform in a syngeneic mouse model of CT26 colorectal cancer using a modified tumour rejection antigen AH1 in combination with anti-PD-1 immune checkpoint inhibitor therapy. AH1 represents one of the best characterized tumour rejection antigens in mice and is derived from the gp70 envelope protein of murine leukaemia virus (MuLV), which is endogenous in the genome of most laboratory mouse strains, including the BALB/c strain used in these studies ^23^. Starting at 11 days post tumour engraftment, mice were treated intratumourally with BCG, anti-PD-1 alone, PeptiBAC-AH1, BCG in combination with anti-PD-1, PeptiBAC-AH1 in combination with anti-PD-1 or saline as a mock-treated group. Once again, the tumour size threshold was set to 450 mm^3^ for defining the responders in each treatment group. Mock, BCG, anti-PD-1 alone and BCG in combination with anti-PD-1 ICI-treated groups showed similar tumour growth characteristics with response rates of 25%, 22%, 25% and 10%, respectively. Interestingly, in contrast to the B16.F10.9/K1 melanoma model, PeptiBAC-AH1 treatment alone did not increase tumour growth control relative to the other groups, with a response rate of 25%. Strikingly, PeptiBAC-AH1 in combination with anti-PD-1-treated animals showed very efficient tumour growth control with an 80% response rate; increasing the response rate for anti-PD-1 therapy from 25% to 80% (Figure 6A). Again, we assessed whether there were any differences in T cell responses between the treatment groups. We saw no significant differences in the numbers of tumour infiltrating CD4^+^ and CD8^+^ T cells between the treatment groups although, interestingly, the number of CD8^+^ T cells in the PeptiBAC-AH1 treated tumours was slightly decreased compared to tumours from other treatment groups. While the number of AH1-specific CD8^+^ T cells was slightly decreased in BCG and BCG in combination with anti-PD-1 ICI-treated tumours when compared to other treatment groups, PeptiBAC-AH1 in combination with anti-PD-1 ICI-treated tumours had significantly increased numbers of AH1-specific CD8^+^ T cells, suggesting a correlation between tumour growth control and the number of AH1-specific CD8^+^ T cells in the TME (Figure 6B, upper panel). Analysis of systemic tumour-specific T cell responses from the spleens of the treated mice showed no significant differences in the number of CD4^+^ and CD8^+^ T cells between groups. However, a significant increase in AH1-specific CD8^+^ T cells was seen in the PeptiBAC-AH1 and PeptiBAC-AH1 in combination with anti-PD-1 ICI-treated mice spleens as compared to spleens from other groups (Figure 6B, lower panel).

**Figure 6.**
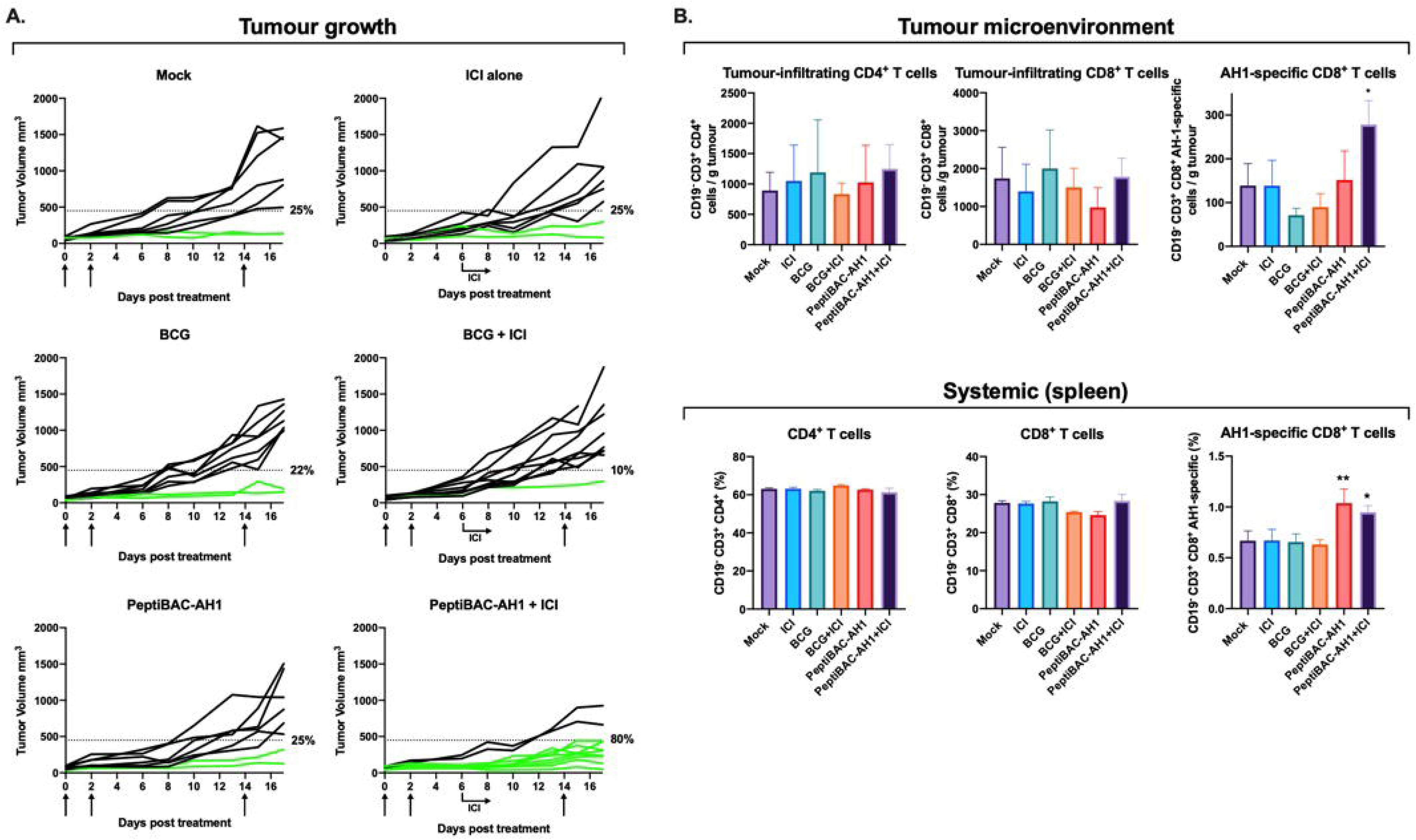
PeptiBAC in combination with anti-PD1 improves tumour growth control compared to either monotherapy and induces systemic tumour-specific CD8^+^ T cell responses and robust infiltration of tumour-specific CD8^+^ T cells into the tumour in a syngeneic mouse model of CT26 colorectal cancer. A) Anti-PD-1 immune checkpoint inhibitor alone (100μg/dose given intraperitoneally three times a week, starting at day 6), BCG alone or in combination with anti-PD-1 immune checkpoint inhibitor and PeptiBAC-AH1 alone or in combination with anti-PD-1 immune checkpoint inhibitor was given intratumourally 11-, 13-, and 25-days post tumour implantation. Individual tumour growth curves for all treatment groups are shown. A threshold of 450 mm^3^ was set to define the percentage of mice responding to the different therapies (dotted line). The percentage of responders in each treatment group is shown on the right side of the dotted line. B) Immunological analysis of tumours and spleens of treated mice. The number of mice in each group was 8-10. Statistical analysis was performed with one-way ANOVA. * p< 0.05, ** p< 0.01.

### Heterologous prime-boost vaccination strategy combining PeptiBAC platform with PeptiCRAd platform improves T cell responses against the coated antigen

Finally, the PeptiBAC-platform was tested in combination with our recently described cancer vaccine platform PeptiCRAd ^14^ (**pepti**de-coated **c**onditionally **r**eplicating **ad**enovirus) using a heterologous prime-boost vaccination strategy. By combining two immunologically distinct platforms coated with the same antigen, we tested whether this heterologous prime-boost approach could enhance T cell-specific immune responses in naïve mice towards the MHC-I restricted epitope presented by both platforms. To this end, we vaccinated naïve C57BL/6JOlaHsd mice with two doses of PeptiBAC-Trp2 or PeptiCRAd-Trp2 as homologous prime-boost controls or with PeptiBAC-Trp2 prime followed by PeptiCRAd-Trp2 boost and PeptiCRAd-Trp2 prime followed by PeptiBAC-Trp2 boost with doses given 14 days apart. 4 days after the boost dose, mice where sacrificed and the spleens were harvested and analysed for the induction of Trp2-specific T cell responses by interferon-gamma ELISPOT. Vaccination with PeptiCRAd-Trp2 homologous prime-boost or PeptiCRAd-Trp2 - PeptiBAC-Trp2 heterologous prime-boost did not induce significant Trp2-specific T cell responses in this vaccination setting. PeptiBAC-Trp2 homologous prime-boost vaccination induced moderate Trp2-specific T cell responses which were markedly enhanced by the PeptiBAC-Trp2 - PeptiCRAd-Trp2 heterologous prime-boost vaccination regimen (Figure 7A). Subsequently, we tested the same approach using the immunodominant epitope of ovalbumin (SIINFEKL), an epitope more immunogenic than Trp2, and assessed the induction of OVA-specific T cell responses again by using the interferon-gamma ELISPOT. Again, the PeptiBAC-OVA - PeptiCRAd-OVA heterologous prime-boost regimen induced significant enhancement of OVA-specific T cell responses compared to PeptiBAC-OVA vaccination (Figure 7B).

**Figure 7.**
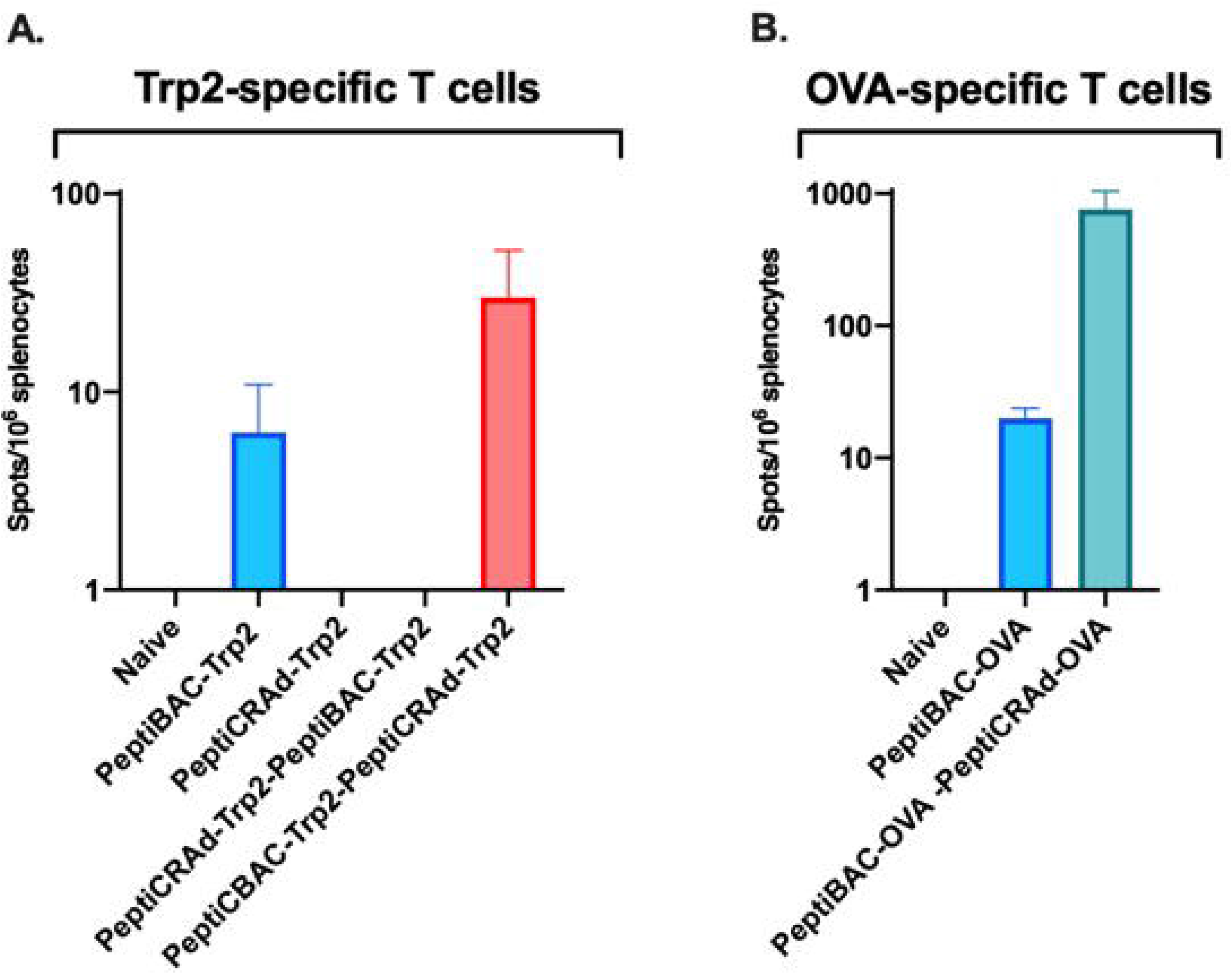
Heterologous prime-boost vaccination with PeptiCRAd platform improves peptide-specific T cell responses elicited by the PeptiBAC platform. A) Naïve C57BL/6JOlaHsd immunocompetent mice were vaccinated subcutaneously with 1×10^9^ VP/dose of PeptiCRAd-Trp2 or 2-8×10^6^ C.F.U/dose of PeptiBAC-Trp2 or saline as a mock-treated group. Prime and boost vaccinations were performed 14 days apart and 4 days after the boost, mice were sacrificed, and spleens were collected for enzyme-linked immunospot (ELISPOT) assay. The number of mice in each vaccination group was 4, and in control group not receiving vaccinations the number of mice was 2. B). Similarly to A, mice were vaccinated with PeptiBAC-OVA or PeptiBAC-OVA followed by PeptiCRAd-OVA booster. The number of mice in each vaccination group was 5.

## Discussion

In this study we have shown that by coating the mycobacterial outer membrane of Bacillus Calmette-Guérin with MHC class I-restricted tumour-associated epitopes, we were able to redirect the immune responses elicited by the bacteria towards the coated peptides. As the attachment moiety for coating the therapeutic peptides onto the mycobacterial outer membrane, we tested both the CPP sequence of the HIV Tat protein fused to the N terminus of the tumour epitopes, and a stretch of 6 lysine residues similarly fused to the N terminus of the tumour epitopes. We have previously shown that the CPP sequence and the polylysine sequence at the N-terminus of the therapeutic peptides do not influence the presentation of the tumour epitopes from these peptides by APCs ^14, 24^. Both attachment moieties were able to efficiently attach therapeutic peptides onto the mycobacterial outer membrane, and BCG coated with an immunodominant epitope derived from chicken ovalbumin (PeptiBAC-OVA) was able to deliver these peptides into APCs followed by efficient processing and presentation by the APCs. The anti-tumour and immune-activating properties of PeptiBAC-OVA was tested in a syngeneic mouse model of B16.OVA melanoma. Although PeptiBAC-OVA induced significant systemic OVA-specific T cell responses, the effect on tumour growth control was moderate at best with the used dose of BCG within the PeptiBAC-OVA. In line with earlier reports ^25, 26^, we did not observe any beneficial effect on tumour growth control by intratumoural treatment with BCG. Interestingly, while PeptiBAC-OVA-treated mice had the longest average survival, we observed a trend towards decreased survival with the BCG-treated group of mice. As poly-lysine sequence also enabled efficient coating of therapeutic peptides onto the mycobacterial outer membrane, we next tested the effects of a higher dose of PeptiBAC with poly-lysine-containing Trp2 epitope (PeptiBAC-Trp2) in combination with immune checkpoint inhibitor (ICI) therapy using an antibody against murine PD-1 in a syngeneic mouse model of B16.F10.9/K1 melanoma. In this model, monotherapy with PeptiBAC-Trp2 induced a clear increase in tumour growth control as compared to Mock, BCG, ICI or BCG + ICI-treated groups. Remarkably, PeptiBAC-Trp2 treatment efficiently sensitized tumours to ICI therapy and the combination therapy group showed a response rate of 70%. In addition to increased tumour growth control, immunological analysis of the treated tumours revealed significant infiltration of CD4^+^, CD8^+^ as well as Trp2-specific CD8^+^ T cells into the TME of the PeptiBAC-Trp2 + ICI-treated mice.

To further evaluate the PeptiBAC platform, we tested the platform in a syngeneic mouse model of CT26 colorectal cancer using a modified tumour rejection antigen AH1 in combination with anti-PD-1 ICI therapy. In this model, although we did not see effects on tumour growth with either monotherapies, the combination of PeptiBAC-AH1 and anti-PD-1 ICI had remarkable synergistic effects, showing a response rate of 80%. In addition, the combo-treated mice showed significantly increased infiltration of AH1-specific CD8^+^ T cells into the TME. Both PeptiBAC-AH1 monotherapy and PeptiBAC-AH1 in combination with anti-PD-1 significantly increased AH1-specific CD8^+^ T cells in spleens as compared to other treatment groups.

Heterologous prime-boost vaccination sequentially using two or more immunologically distinct platforms to deliver the antigen(s) has previously been tested in both infectious disease and cancer settings ^27–31^, and has shown to be able to induce enhanced T cell responses against the antigen as compared to homologous prime-boost vaccination. Also, BCG has previously been used as a component in heterologous prime-boost settings ^32–34^. Here we set out to test whether the PeptiBAC platform could be used as a component of a heterologous prime-boost vaccination setting together with another peptide-based cancer vaccine platform using oncolytic adenoviruses, called PeptiCRAd. Interestingly, we saw enhanced antigen-specific T cell responses as compared to homologous prime-boost vaccination with PeptiBAC only when PeptiBAC was used as a priming vaccine and PeptiCRAd as a booster vaccine.

In addition to CIS, BCG is the preferred treatment for high-risk non-muscle-invasive bladder cancer (NMIBC) and an option for intermediate-risk NMIBC ^35^. Recently, the US Food and Drug Administration approved an immune checkpoint inhibitor against PD-1 (pembrolizumab) to treat patients with BCG-unresponsive, high-risk, NMIBC with carcinoma in situ with or without papillary tumours who are ineligible for, or have elected not to undergo cystectomy ^36^. In addition, a recent phase III trial that evaluated a novel intravesical therapy, nadofaragene firadenovec (a non-replicating adenovirus vector expressing human IFNα2b) in 151 patients with BCG-unresponsive NMIBC reported that more than half of the patients achieved a complete response, of whom almost half maintained complete response at 12 months ^37^. It is intriguing to hypothesize, in light of the data presented here, that using PeptiBAC with tumour specific (neo)antigens identified from bladder cancer to treat NMIBC could increase the response rate of BCG therapy, and in addition, if used in combination with pembrolizumab, could have significant improvements over outcomes achieved with BCG or pembrolizumab as monotherapies. Nadofaragene firadenovec is compatible with the PeptiCRAd cancer vaccine platform and could be tested as part of the PeptiCRAd platform together with prior therapy with PeptiBAC as a heterologous prime-boost cancer vaccine immunotherapy. Compared to various other immunotherapy approaches, the PeptiBAC platform is highly adaptable and can be quickly coated with a patient’s unique set of tumour-specific antigens, a prerequisite for personalized cancer immunotherapy. Most importantly, this platform could be transferred into the clinical setting very fast, since the backbone of the platform, the BCG vaccine, is already FDA/EMEA approved for cancer immunotherapy for bladder cancer and melanoma.

In addition to being used as a cancer immunotherapy, BCG is the only vaccine used in infants and neonates to prevent tuberculous meningitis and disseminated tuberculosis ^38^. Remarkably, in addition to its specific effect against tuberculosis, the BCG vaccine has beneficial non-specific (off-target) effects on the immune system that protect against a wide range of other infections, including bacteria like *Staphylococcus aureus*, fungi like *Candida albicans* and viruses like the yellow fever virus ^39, 40^. Recent studies have suggested that countries that mandate BCG vaccination for the population have a lower number of infections and a reduced mortality from COVID-19 ^41^. Based on these data, it has been hypothesized that BCG vaccination might be a potent preventive measure against SARS-CoV-2 infection and/or may reduce COVID-19 disease severity. Currently, there are at least 9 clinical studies ongoing to determine the effect of BCG vaccination on outcomes from COVID-19. However, the efficacy of the BCG vaccine to provide protection against COVID-19 might be significantly improved by enhancing the SARS-CoV-2-specific cellular immune responses elicited by the BCG vaccine by the use of PeptiBAC platform with SARS-CoV-2-specific antigens.

## Acknowledgements

E.Y acknowledges the Academy of Finland (project N° 1317206) and HiLIFE Proof-of-Concept grant (project N° 115422). V.C. acknowledges the European Research Council under the Horizon 2020 framework (https://erc.europa.eu), ERC-consolidator Grant (Agreement N° 681219), Jane and Aatos Erkko Foundation (Project N° 4705796), HiLIFE Fellow (project N° 797011004), Cancer Finnish Foundation (project N° 4706116), Magnus Ehrnrooth Foundation (project N° 4706235), Academy of Finland and Digital Precision Cancer Medicine Flagship iCAN.

## References

1. Kamat, A. M.; Colombel, M.; Sundi, D.; Lamm, D.; Boehle, A.; Brausi, M.; Buckley, R.; Persad, R.; Palou, J.; Soloway, M.; Witjes, J. A., BCG-unresponsive non-muscle-invasive bladder cancer: recommendations from the IBCG. Nat Rev Urol 2017, 14 (4), 244–255.

2. Tse, J.; Singla, N.; Ghandour, R.; Lotan, Y.; Margulis, V., Current advances in BCG-unresponsive non-muscle invasive bladder cancer. Expert Opin Investig Drugs 2019, 28 (9), 757–770.

3. Morton, D.; Eilber, F. R.; Malmgren, R. A.; Wood, W. C., Immunological factors which influence response to immunotherapy in malignant melanoma. Surgery 1970, 68 (1), 158–63; discussion 163-4.

4. Morton, D. L.; Eilber, F. R.; Holmes, E. C.; Hunt, J. S.; Ketcham, A. S.; Silverstein, M. J.; Sparks, F. C., BCG immunotherapy of malignant melanoma: summary of a seven-year experience. Ann Surg 1974, 180 (4), 635–43.

5. Coit, D. G.; Thompson, J. A.; Algazi, A.; Andtbacka, R.; Bichakjian, C. K.; Carson, W. E., 3rd; Daniels, G. A.; DiMaio, D.; Ernstoff, M.; Fields, R. C.; Fleming, M. D.; Gonzalez, R.; Guild, V.; Halpern, A. C.; Hodi, F. S., Jr.; Joseph, R. W.; Lange, J. R.; Martini, M. C.; Materin, M. A.; Olszanski, A. J.; Ross, M. I.; Salama, A. K.; Skitzki, J.; Sosman, J.; Swetter, S. M.; Tanabe, K. K.; Torres-Roca, J. F.; Trisal, V.; Urist, M. M.; McMillian, N.; Engh, A., Melanoma, Version 2.2016, NCCN Clinical Practice Guidelines in Oncology. J Natl Compr Canc Netw 2016, 14 (4), 450–73.

6. Kibbi, N.; Ariyan, S.; Faries, M.; Choi, J. N., Treatment of in-transit melanoma with intralesional bacillus Calmette-Guérin (BCG) and topical imiquimod 5% cream: a report of 3 cases. J Immunother 2015, 38 (9), 371–5.

7. Kidner, T. B.; Morton, D. L.; Lee, D. J.; Hoban, M.; Foshag, L. J.; Turner, R. R.; Faries, M. B., Combined intralesional Bacille Calmette-Guérin (BCG) and topical imiquimod for in-transit melanoma. J Immunother 2012, 35 (9), 716–20.

8. Redelman-Sidi, G.; Glickman, M. S.; Bochner, B. H., The mechanism of action of BCG therapy for bladder cancer--a current perspective. Nat Rev Urol 2014, 11 (3), 153–62.

9. Yang, J.; Jones, M. S.; Ramos, R. I.; Chan, A. A.; Lee, A. F.; Foshag, L. J.; Sieling, P. A.; Faries, M. B.; Lee, D. J., Insights into Local Tumor Microenvironment Immune Factors Associated with Regression of Cutaneous Melanoma Metastases by Mycobacterium bovis Bacille Calmette-Guérin. Front Oncol 2017, 7, 61.

10. Antonelli, A. C.; Binyamin, A.; Hohl, T. M.; Glickman, M. S.; Redelman-Sidi, G., Bacterial immunotherapy for cancer induces CD4-dependent tumor-specific immunity through tumor-intrinsic interferon-γ signaling. Proc Natl Acad Sci U S A 2020, 117 (31), 18627–18637.

11. Schoenfeld, A. J.; Hellmann, M. D., Acquired Resistance to Immune Checkpoint Inhibitors. Cancer Cell 2020, 37 (4), 443–455.

12. Ku, G. Y.; Yuan, J.; Page, D. B.; Schroeder, S. E.; Panageas, K. S.; Carvajal, R. D.; Chapman, P. B.; Schwartz, G. K.; Allison, J. P.; Wolchok, J. D., Single-institution experience with ipilimumab in advanced melanoma patients in the compassionate use setting: lymphocyte count after 2 doses correlates with survival. Cancer 2010, 116 (7), 1767–75.

13. Yuan, J.; Adamow, M.; Ginsberg, B. A.; Rasalan, T. S.; Ritter, E.; Gallardo, H. F.; Xu, Y.; Pogoriler, E.; Terzulli, S. L.; Kuk, D.; Panageas, K. S.; Ritter, G.; Sznol, M.; Halaban, R.; Jungbluth, A. A.; Allison, J. P.; Old, L. J.; Wolchok, J. D.; Gnjatic, S., Integrated NY-ESO-1 antibody and CD8+ T-cell responses correlate with clinical benefit in advanced melanoma patients treated with ipilimumab. Proc Natl Acad Sci U S A 2011, 108 (40), 16723–8.

14. Capasso, C.; Hirvinen, M.; Garofalo, M.; Romaniuk, D.; Kuryk, L.; Sarvela, T.; Vitale, A.; Antopolsky, M.; Magarkar, A.; Viitala, T.; Suutari, T.; Bunker, A.; Yliperttula, M.; Urtti, A.; Cerullo, V., Oncolytic adenoviruses coated with MHC-I tumor epitopes increase the antitumor immunity and efficacy against melanoma. Oncoimmunology 2016, 5 (4), e1105429.

15. Vordermeier, H. M.; Rhodes, S. G.; Dean, G.; Goonetilleke, N.; Huygen, K.; Hill, A. V.; Hewinson, R. G.; Gilbert, S. C., Cellular immune responses induced in cattle by heterologous prime-boost vaccination using recombinant viruses and bacille Calmette-Guérin. Immunology 2004, 112 (3), 461–70.

16. Evans, R. K.; Nawrocki, D. K.; Isopi, L. A.; Williams, D. M.; Casimiro, D. R.; Chin, S.; Chen, M.; Zhu, D. M.; Shiver, J. W.; Volkin, D. B., Development of stable liquid formulations for adenovirus-based vaccines. J Pharm Sci 2004, 93 (10), 2458–75.

17. Jordan, K. R.; McMahan, R. H.; Kemmler, C. B.; Kappler, J. W.; Slansky, J. E., Peptide vaccines prevent tumor growth by activating T cells that respond to native tumor antigens. Proc Natl Acad Sci U S A 2010, 107 (10), 4652–7.

18. Bansal-Mutalik, R.; Nikaido, H., Mycobacterial outer membrane is a lipid bilayer and the inner membrane is unusually rich in diacyl phosphatidylinositol dimannosides. Proc Natl Acad Sci U S A 2014, 111 (13), 4958–63.

19. Kristensen, S.; Tian, Y.; Klegerman, M. E.; Groves, M. J., Origins of BCG surface charge: effect of ionic strength and chemical modifications on zeta potential of Mycobacterium bovis BCG, Tice substrain, cells. Microbios 1992, 70 (284-285), 185–98.

20. Zhang, A.; Groves, M. J.; Klegerman, M. E., The surface charge of cells of Mycobacterium bovis BCG vaccine, Tice substrain. Microbios 1988, 53 (216-217), 191–5.

21. Moore, M. W.; Carbone, F. R.; Bevan, M. J., Introduction of soluble protein into the class I pathway of antigen processing and presentation. Cell 1988, 54 (6), 777–85.

22. Porgador, A.; Feldman, M.; Eisenbach, L., H-2Kb transfection of B16 melanoma cells results in reduced tumourigenicity and metastatic competence. J Immunogenet 1989, 16 (4-5), 291–303.

23. Jenkins, N. A.; Copeland, N. G.; Taylor, B. A.; Lee, B. K., Organization, distribution, and stability of endogenous ecotropic murine leukemia virus DNA sequences in chromosomes of Mus musculus. J Virol 1982, 43 (1), 26–36.

24. Ylösmäki, E.; Malorzo, C.; Capasso, C.; Honkasalo, O.; Fusciello, M.; Martins, B.; Ylösmäki, L.; Louna, A.; Feola, S.; Paavilainen, H.; Peltonen, K.; Hukkanen, V.; Viitala, T.; Cerullo, V., Personalized Cancer Vaccine Platform for Clinically Relevant Oncolytic Enveloped. Mol Ther 2018, 26 (9), 2315–2325 LID - S1525-0016(18)30267-3 [pii] LID - 10.1016/j.ymthe.2018.06.008 [doi].

25. Kreider, J. W.; Bartlett, G. L.; Purnell, D. M., Inconsistent response of B16 melanoma to BCG immunotherapy. J Natl Cancer Inst 1976, 56 (4), 803–10.

26. Piessens, W. F.; Lachapelle, F. L.; Legros, N.; Heuson, J. C., Facilitation of rat mammary tumour growth by BCG. Nature 1970, 228 (5277), 1210–1.

27. Aitken, A. S.; Roy, D. G.; Martin, N. T.; Sad, S.; Bell, J. C.; Bourgeois-Daigneault, M. C., Brief Communication; A Heterologous Oncolytic Bacteria-Virus Prime-Boost Approach for Anticancer Vaccination in Mice. J Immunother 2018, 41 (3), 125–129.

28. Bridle, B. W.; Boudreau, J. E.; Lichty, B. D.; Brunellière, J.; Stephenson, K.; Koshy, S.; Bramson, J. L.; Wan, Y., Vesicular stomatitis virus as a novel cancer vaccine vector to prime antitumor immunity amenable to rapid boosting with adenovirus. Mol Ther 2009, 17 (10), 1814–21.

29. Hu, S. L.; Abrams, K.; Barber, G. N.; Moran, P.; Zarling, J. M.; Langlois, A. J.; Kuller, L.; Morton, W. R.; Benveniste, R. E., Protection of macaques against SIV infection by subunit vaccines of SIV envelope glycoprotein gp160. Science 1992, 255 (5043), 456–9.

30. Hu, S. L.; Klaniecki, J.; Dykers, T.; Sridhar, P.; Travis, B. M., Neutralizing antibodies against HIV-1 BRU and SF2 isolates generated in mice immunized with recombinant vaccinia virus expressing HIV-1 (BRU) envelope glycoproteins and boosted with homologous gp160. AIDS Res Hum Retroviruses 1991, 7 (7), 615–20.

31. Pol, J. G.; Acuna, S. A.; Yadollahi, B.; Tang, N.; Stephenson, K. B.; Atherton, M. J.; Hanwell, D.; El-Warrak, A.; Goldstein, A.; Moloo, B.; Turner, P. V.; Lopez, R.; LaFrance, S.; Evelegh, C.; Denisova, G.; Parsons, R.; Millar, J.; Stoll, G.; Martin, C. G.; Pomoransky, J.; Breitbach, C. J.; Bramson, J. L.; Bell, J. C.; Wan, Y.; Stojdl, D. F.; Lichty, B. D.; McCart, J. A., Preclinical evaluation of a MAGE-A3 vaccination utilizing the oncolytic Maraba virus currently in first-in-human trials. Oncoimmunology 2019, 8 (1), e1512329.

32. Li, W.; Li, M.; Deng, G.; Zhao, L.; Liu, X.; Wang, Y., Prime-boost vaccination with Bacillus Calmette Guerin and a recombinant adenovirus co-expressing CFP10, ESAT6, Ag85A and Ag85B of Mycobacterium tuberculosis induces robust antigen-specific immune responses in mice. Mol Med Rep 2015, 12 (2), 3073–80.

33. Magalhaes, I.; Sizemore, D. R.; Ahmed, R. K.; Mueller, S.; Wehlin, L.; Scanga, C.; Weichold, F.; Schirru, G.; Pau, M. G.; Goudsmit, J.; Kühlmann-Berenzon, S.; Spångberg, M.; Andersson, J.; Gaines, H.; Thorstensson, R.; Skeiky, Y. A.; Sadoff, J.; Maeurer, M., rBCG induces strong antigen-specific T cell responses in rhesus macaques in a prime-boost setting with an adenovirus 35 tuberculosis vaccine vector. PLoS One 2008, 3 (11), e3790.

34. Xu, Y.; Yang, E.; Wang, J.; Li, R.; Li, G.; Liu, G.; Song, N.; Huang, Q.; Kong, C.; Wang, H., Prime-boost bacillus Calmette-Guérin vaccination with lentivirus-vectored and DNA-based vaccines expressing antigens Ag85B and Rv3425 improves protective efficacy against Mycobacterium tuberculosis in mice. Immunology 2014, 143 (2), 277–86.

35. Lenis, A. T.; Lec, P. M.; Chamie, K.; Mshs, M. D., Bladder Cancer: A Review. Jama 2020, 324 (19), 1980–1991.

36. Gill, J.; Prasad, V., Pembrolizumab for Non-Muscle-Invasive Bladder Cancer-A Costly Therapy in Search of Evidence. JAMA Oncol 2020.

37. Boorjian, S. A.; Alemozaffar, M.; Konety, B. R.; Shore, N. D.; Gomella, L. G.; Kamat, A. M.; Bivalacqua, T. J.; Montgomery, J. S.; Lerner, S. P.; Busby, J. E.; Poch, M.; Crispen, P. L.; Steinberg, G. D.; Schuckman, A. K.; Downs, T. M.; Svatek, R. S.; Mashni, J., Jr.; Lane, B. R.; Guzzo, T. J.; Bratslavsky, G.; Karsh, L. I.; Woods, M. E.; Brown, G.; Canter, D.; Luchey, A.; Lotan, Y.; Krupski, T.; Inman, B. A.; Williams, M. B.; Cookson, M. S.; Keegan, K. A.; Andriole, G. L., Jr.; Sankin, A. I.; Boyd, A.; O’Donnell, M. A.; Sawutz, D.; Philipson, R.; Coll, R.; Narayan, V. M.; Treasure, F. P.; Yla-Herttuala, S.; Parker, N. R.; Dinney, C. P. N., Intravesical nadofaragene firadenovec gene therapy for BCG-unresponsive non-muscle-invasive bladder cancer: a single-arm, open-label, repeat-dose clinical trial. Lancet Oncol 2021, 22 (1), 107–117.

38. Tran, V.; Liu, J.; Behr, M. A., BCG Vaccines. Microbiol Spectr 2014, 2 (1), Mgm2-0028-2013.

39. Arts, R. J. W.; Moorlag, S.; Novakovic, B.; Li, Y.; Wang, S. Y.; Oosting, M.; Kumar, V.; Xavier, R. J.; Wijmenga, C.; Joosten, L. A. B.; Reusken, C.; Benn, C. S.; Aaby, P.; Koopmans, M. P.; Stunnenberg, H. G.; van Crevel, R.; Netea, M. G., BCG Vaccination Protects against Experimental Viral Infection in Humans through the Induction of Cytokines Associated with Trained Immunity. Cell Host Microbe 2018, 23 (1), 89–100.e5.

40. Kleinnijenhuis, J.; Quintin, J.; Preijers, F.; Benn, C. S.; Joosten, L. A.; Jacobs, C.; van Loenhout, J.; Xavier, R. J.; Aaby, P.; van der Meer, J. W.; van Crevel, R.; Netea, M. G., Long-lasting effects of BCG vaccination on both heterologous Th1/Th17 responses and innate trained immunity. J Innate Immun 2014, 6 (2), 152–8.

41. Gursel, M.; Gursel, I., Is global BCG vaccination-induced trained immunity relevant to the progression of SARS-CoV-2 pandemic? Allergy 2020, 75 (7), 1815–1819.

